# Finger representation in the cortex of the congenitally blind

**DOI:** 10.1101/2021.03.16.435392

**Authors:** D.B. Wesselink, S. Kikkert, H. Bridge, T.R. Makin

**Author notes:** **Corresponding author**: Tamar Makin, Institute of Cognitive Neuroscience, University College London, London WC1N 3AZ;).

## Abstract

Hand representation in the primary somatosensory cortex (S1) is thought to be shaped by experience. Individuals with congenital blindness rely on their sense of touch for completing daily tasks that in sighted people would be informed by vision, and possess superior tactile acuity. It has therefore been proposed that their S1 hand representation should differ from that of sighted individuals. Alternatively, it has been proposed that the improved tactile acuity in blind individuals is due to cross-modal plasticity, when regions in the occipital and temporal cortex are typically used for processing vision become activated by touch. We probed finger representation using psychophysics and 7T fMRI (1 mm^3^ resolution) in three individuals with bilateral anophthalmia, a rare condition in which both eyes fail to develop, as well as sighted controls. Despite anophthalmic individuals’ increased reliance on touch and superior tactile acuity, we found no evidence that they had more pronounced hand representation in S1. This is in line with recent research highlighting the stability of early sensory cortex, despite altered sensorimotor experience in adulthood. Unlike sighted controls, anophthalmic individuals activated the left human middle temporal complex (hMT+) during finger movement. This area did not express any hallmark of typical sensorimotor organisation, suggesting this and previously reported activity does not indicate low-level sensorimotor hand representation. However, left hMT+ contained some single finger information, beyond that found in sighted controls. This latter finding suggests that when the developmentally flexible area hMT+ is unaffected by retinal input, it can acquire novel cross-modal processes, which are potentially unrelated to the area’s function in sighted people. As such, our findings highlight the opportunity for other organising principles, beyond domain specific plasticity, in shaping cross-modal reorganisation.

## Introduction

To successfully interact with their environment, individuals with congenital blindness are forced to rely on other senses, such as touch. Congruently, both congenitally and early blind people show superior performance on a variety of tactile tasks (Goldreich and Kanics 2003, Van Boven et al. 2000, Norman and Bartholomew 2011, though see Alary et al. 2009). Given previous evidence for a relationship between tactile ability and the recruitment of the somatosensory system, particularly the primary somatosensory cortex (S1; e.g. Recanzone et al. 1992), it is not surprising that researchers have hypothesised that S1 hand representation of blind individuals should be altered from that of the sighted. Blind individuals’ fingers, in particular the reading finger(s) of Braille readers, have been reported to be represented in a more pronounced manner in S1, in terms of activity level or spatial spread (Pascual-Leone and Torres 1993, Sterr et al. 1998) but see (Gizewski et al. 2003, Burton et al. 2004); see Discussion for full details]. This is consistent with the suggestion that following increased sensory demands on our fingers, such as by learning to play an instrument (Elbert et al. 1995), or mere repeated exposure to tactile stimuli (Recanzone et al. 1992, Jenkins et al. 1990, Gindrat et al. 2015), cortical magnification becomes more pronounced. Following these findings, it has been proposed that the increased tactile perceptual abilities documented in blind individuals could be caused by different finger representation in S1 (e.g. Van Boven et al. 2000, Lissek et al. 2009).

Other studies have suggested that the increased tactile acuity found in blind individuals may be facilitated, in part, by the (repurposed) ‘visual’ cortex. Cross-modal cortical reorganisation is commonly observed in traditionally visual areas as over-excitability during non-visual tasks (Merabet and Pascual-Leone 2010, Bavelier and Neville 2002). For example, previous research has shown that early blind individuals activate both the primary visual cortex (V1; (Sadato et al. 1996) and occipitotemporal cortex (Burton et al. 2002, Sadato et al. 2004) during Braille reading. Yet, the exact function of emergent processes in the ‘visual’ cortex may not be comparable to that of S1. For example, V1 appears to provide linguistic rather than strictly sensory support during Braille reading (Hamilton et al. 2000, Bedny et al. 2011)]. Regardless, the initial findings led to suggestions that cross-modal reorganisation could support the heightened behavioural needs for touch processing in blind individuals by enabling new functions. Since then, this interpretation has been challenged. Instead, it has been suggested that the specific computations performed by a repurposed area remain similar following cross-modal reorganisation (domain specificity). As such, the only difference between occipital processes of blind and sighted individuals is their input modality (Pascual-Leone and Hamilton 2001, see also Mahon et al. 2009). For example, in the visual cortex of the sighted, the extrastriate body area has been implicated with visual processing of body parts (Downing et al. 2001; Kontaris, Wiggett, and Downing 2009). Accordingly, in blind individuals, this body-selective region is specifically activated during recognition of body parts and body postures through touch (Kitada et al. 2014) or auditory cues (Striem-Amit and Amedi 2014). This and other related findings suggest that cross-modal reorganisation is more restricted than initially thought, with only limited capacity for reassignment of function.

The framework of domain specificity has been highly productive (Ricciardi et al. 2014, Maidenbaum et al. 2014, Kupers and Ptito 2014) and many findings have accumulated supporting the principle for medium to high-level functions (e.g. Striem-Amit et al. 2012, Dormal et al. 2016, Peelen et al. 2013, Amedi et al. 2007, Amedi et al. 2010, Wolbers et al. 2011, Kitada et al. 2014). Yet, restricting the scope for functional reassignment of resources is seemingly at odds with some evidence for emergent low-level sensory processing in the occipitotemporal cortex. Watkins et al. (2013) studied individuals with bilateral anophthalmia, a rare condition in which both eyes completely fail to develop. Unlike other forms of early blindness, anophthalmia guarantees the visual cortex has never been influenced by retinal input, not even prenatally. The authors observed structured activations by pure tones (tonotopy) in the occipitotemporal cortex, corresponding to motion-sensitive area hMT+ (human middle temporal complex; see Huber et al. (2019) for a replication in early blind participants). In other words, hMT+ appears to perform an early (low-level) sensory function, independently from known visual processes in the sighted, or at least appears to have extended its original function beyond a cross-modal input shift (see Shiell 2014). This finding may contradict the framework of domain specificity, but it does not demand developmental anarchy. A moderate account would be that rigid functional domains (i.e. domain specificity) develop when the brain’s coarse proto-organisation (Arcaro and Livingstone 2017, Krubitzer and Prescott 2018) is supplied with typical inputs, irrespective of which modality they are in. Completely different inputs, however, may result in domains that are unrecognizably different, in both function and organisational structure (Bedny 2017).

In the current study, we interrogated the low-level somatosensory processing in the same anophthalmic population mentioned above. We focused on the representation of the fingers, which, especially in the human hand area of S1, shows strong selectivity (Kurth et al. 1998, Sanchez-Panchuelo et al. 2010), as well as an idiosyncratic pattern of inter-finger overlap. These properties are shared by a range of low-to-medium-level sensorimotor areas (Berlot et al. 2019, van der Zwaag et al. 2013, Zeharia et al. 2015). Using 7T fMRI, it is possible to identify these characteristic finger maps in humans, with high inter- and intra-subject reliability (Kolasinski et al. 2016a, Ejaz et al. 2015), providing opportunities for studying specialised case studies (Kikkert et al. 2016, Dempsey-Jones et al. 2019b). We wished to determine whether the anophthalmic individuals’ S1 shows stronger finger representation, in terms of interfinger selectivity and overlap. We also examined their visual cortex (and area hMT+ in particular) to determine whether it supports low-level finger processing.

We focused on hMT+ because it has previously been shown to have undergone low-level functional reorganisation in congenitally blind or blindsighted individuals (Watkins et al. 2013, Ajina and Bridge 2018). Indeed, some have suggested that due to its connectome and in particular its direct connections with sub-cortical structures (Gaglianese et al. 2015), hMT+ resembles a primary area (Ajina et al. 2015a). By effectively being able to bypass V1 inputs, hMT+ can develop an independent functional organisation. As such, this area may be awarded more developmental flexibility (see Ajina et al. (2015b) for structural plasticity) than higher-order visual areas further down-stream. According to this account, functional plasticity can occur here more-or-less unconstrained by the occipital cortex’ organisational template, potentially even allowing hMT+ to develop novel finger representation, organised in a different manner than those found in S1.

We found that despite superior tactile perceptual abilities and a lifetime of increased reliance on touch for typically visual tasks, S1 finger representation was not more pronounced in anophthalmic individuals, relative to controls. Additionally, finger-specific information was detectible in the left hMT+ area of anophthalmic individuals, although the organisational representational structure in occipitotemporal cortex differed from hand somatotopy. Our findings show that, in exceptional circumstances, the visual cortex retains some ability to acquire functional organisation which might not be native to the region, suggesting the prevalent principle of domain specificity does not bridle the development of all sensory areas to the same extent.

## Results

### Increased tactile acuity in anophthalmic individuals

First, we measured tactile acuity using a psychophysical task: participants were asked to judge whether gratings of varying fineness were placed horizontally or vertically on their fingertips. This allowed us to construct a psychometric (Weibull) function per finger and participant to determine the tactile threshold (defined at 82% correct performance; see Methods). The average tactile threshold (across the three fingers tested) was significantly lower in the anophthalmic individuals compared with sighted participants (U=15, p=0.01; see Figure 1), indicating greater tactile acuity. This is in accordance with previous reports showing superior tactile acuity in the congenitally blind (Van Boven et al. 2000, Norman and Bartholomew 2011).

**Figure 1:**
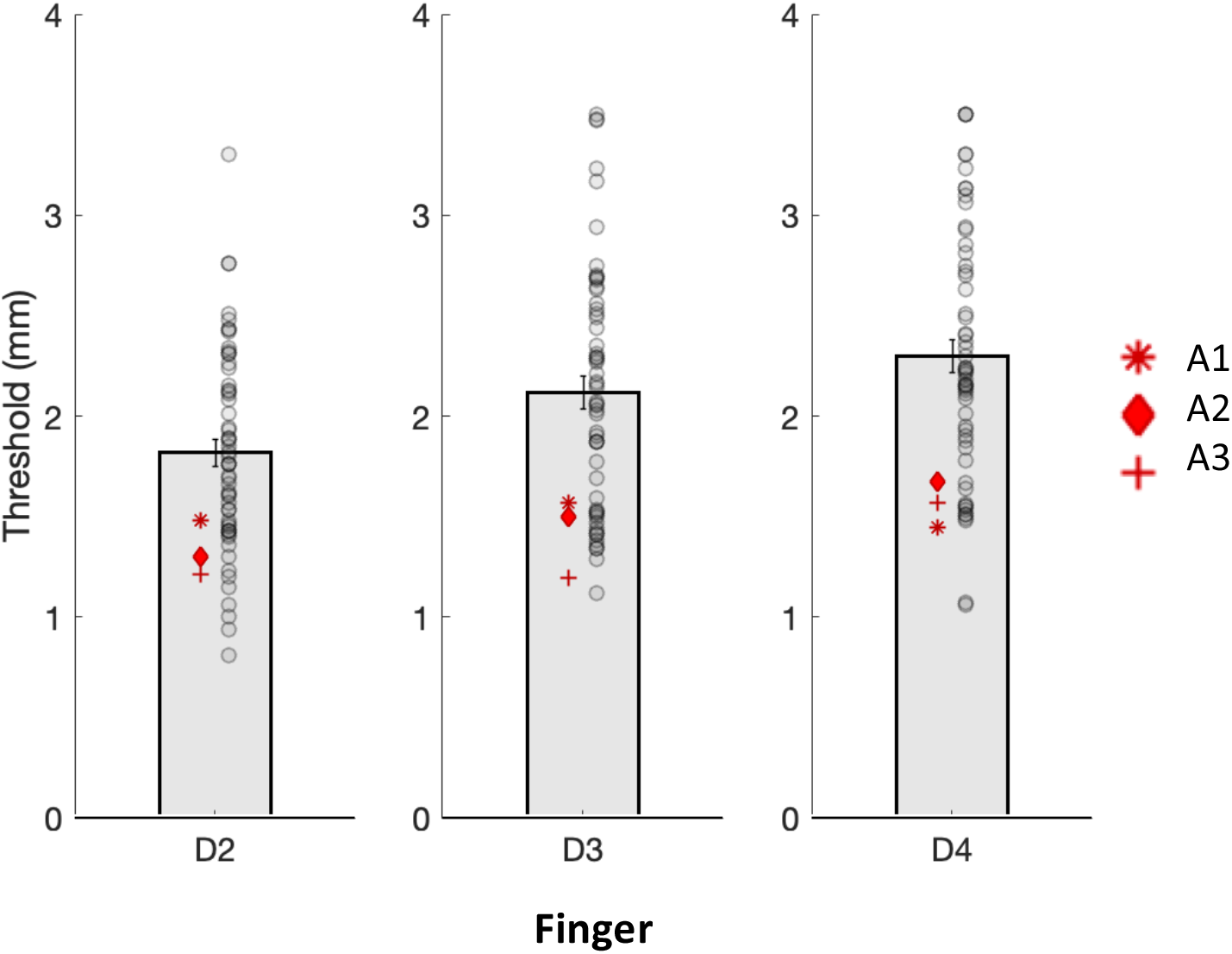
Tactile thresholds in anophthalmic individuals (n=3; red symbols) and sighted controls (n=57; grey circles) for index (D2), middle (D3), and ring finger (D4). On average, the anophthalmic individuals (A1-3) showed greater tactile acuity (lower thresholds) then the sighted controls. Bars shows control mean; error bars indicate the standard errors.

### Hand representations in S1 is similar in anophthalmic individuals and sighted controls

To explore whether S1 would reflect these improved tactile abilities, we first interrogated finger representations using fMRI. All participants performed single-finger movements in a blocked design. As expected, for all participants, there was clear finger selectivity in S1, defined by contrasting each finger’s movement activity against the mean activity for all other fingers. Selectivity maps were qualitatively similar between the anophthalmic individuals and controls (see Figures 2A and S1). Classical univariate analysis did not indicate hand representation was more pronounced in the anophthalmic individuals: neither average activity during finger movement (U=23, p=.376) nor the spatial spread of the hand map, i.e. the number of voxels that showed finger selectivity (U=23, p=.376), were different between groups. As a follow-up, however, when we separately tested whether the spread of each individual finger’s representation had increased, representation of the thumb (D1) of the anophthalmic participants was significantly more widespread (U=27, p=.036, one-sided), but not that of other fingers (p’s > .497). To further quantify hand representation, we calculated multivariate measures, previously suggested to reflect habitual behaviour in typical populations (Ejaz et al. 2015). Specifically, we performed representational similarity analysis (RSA; see Methods) on the activity patterns in S1 for each finger to calculate a value (i.e. dissimilarity value) for each finger pair indicating how distinctly the fingers activated S1. Mean dissimilarity indicates finger separability. All finger pairs’ dissimilarity values together define a representational structure that is a low-dimensional summary of hand representation. Neither measure was greater for the anophthalmic individuals than for the sighted controls (separability: U=12.5, p=.303; representational structure: U=17, p=.921). This suggest the greater spatial spread for D1 activity, reported above, is not reflected in the multivariate hand representation.

**Figure 2:**
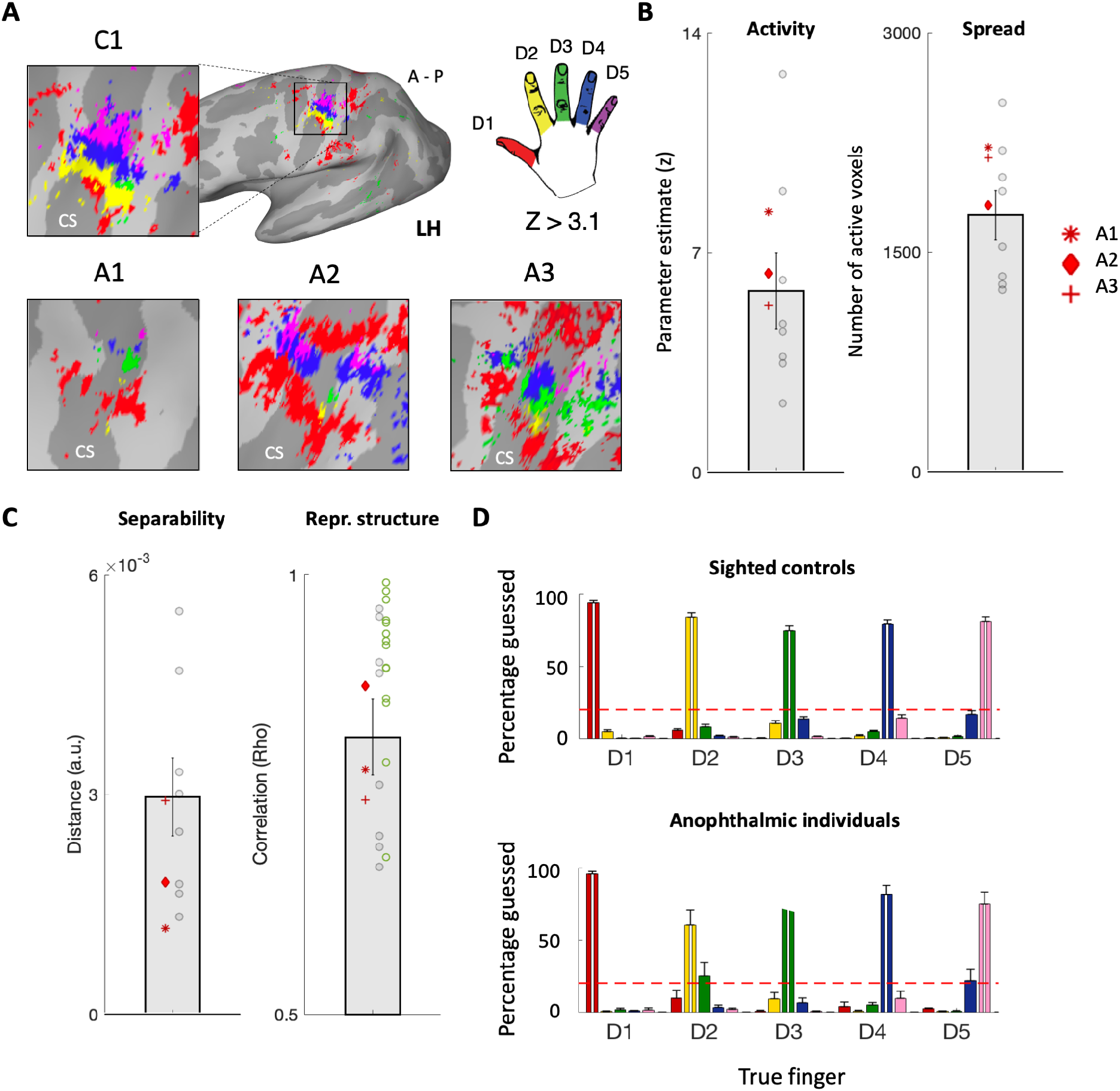
Hand representation in primary somatosensory cortex (S1). **A)** Finger maps for three anophthalmic individuals and a representative sighted control in S1. The sighted control (C1) had median (4^th^/8) average classification performance on fingers (88.1%; see D). Colours indicate selectivity (Z>3.1) for each finger relative to the average of all other fingers (see hand legend for colour code). Each of the anophthalmic participants presented distinct clusters showing selectivity to each of the five fingers, in a typical lateral to medial order. Note that the double D1 representation is commonly observed in sighted controls (e.g. Figure S1 in Sanders et al. (2019); for finger maps of all controls see Figure S1). **B)** Univariate proxy measures of hand representation are not higher in anophthalmic individuals than sighted controls. Activity relates to the average parameter estimate (z) for all fingers versus rest; Spread relates to the number of voxels surviving correction for multiple comparisons within the S1 area. **C)** Multivariate proxy measures are not higher in anophthalmic individuals than sighted controls. Separability and representational structure are two outcome measures of representational similarity analysis. Separability indicates average dissimilarity (representational individuality of each finger pair). Representational structure indicates correlation between the inter-finger representation for each individual subject and to the mean group representation in controls. Supplementary pattern structure data obtained from a separate dataset are indicated in green (n=15). **D)** Average classification performance across participants for sighted controls (top) and anophthalmic individuals (bottom). Chance classifier performance is at 20% and perfect performance is at 100%. For each true finger (highlighted with a white vertical line), few trials were misclassified (neighbouring bars; colours as in A). Misclassifications tended to be more prominent for the neighbouring digits (e.g. D3 was most commonly confused with D2 and D4) in both sighted and anophthalmic groups. Other symbols as in Figure 1.

In short, in all whole-hand measures, the anophthalmic individuals did not significantly differ from controls (Figure 2B-C). We followed up on our original group comparisons using a Bayesian one-tailed t-test, designed to produce conclusive evidence in favour of the null (i.e. the blind group not showing greater finger representation relative to controls). Both the average activity (BF=.659) and the spatial spread of the hand map (BF=1.08) produced inconclusive results, but there was substantial evidence that the anophthalmic individuals did not show greater separability than sighted controls (BF=.314). A Bayesian t-test on the representational structure was inconclusive (BF=.454). Together, these results suggest that the enhanced tactile abilities commonly observed in blind people may not result from more pronounced finger representation in S1. Yet, the inconclusive results for some of our Bayesian t-tests may have resulted from our small sample size. A key advantage of our multivariate measure for representational structure is its high consistency both within and across participants (Ejaz et al. 2015, Walther et al. 2016), particularly when compared to traditional finger selectivity analyses using univariate measures. To further ameliorate the concern of small sample size, we compared the representational structure from the anophthalmic individuals to a relatively large distribution of sighted controls, by combining data from this study with that of another study with similar methodology (15 additional control participants, (Sanders et al. 2019), see Methods). When repeating the statistical comparison with the larger control group, we again found that the representational structure was not different between the anophthalmic individuals and sighted controls; U=24, p=.199). A Bayesian t-test confirmed that there was substantial evidence (BF=.285) to support that the anophthalmic individuals do not show greater (or less noisy) hand representation than the sighted controls.

Finally, as an additional measure of finger selectivity, we trained a basic artificial neural network on the unthresholded multivariate patterns in S1 to classify individual finger movements (leave-one-out classification; see Methods). Although the classifier tells us little about the structure of S1 hand representation, it is a sensitive indicator of finger-specific information in the area. As expected, the classifier was able to correctly classify the five fingers (chance level 20%; created using finger label permutation) with high accuracy for both anophthalmic individuals (mean=79.1%) and sighted controls (mean=82.6%; Figure 2D). When comparing the average classification accuracy between the two groups, the anophthalmic individuals did not show greater classification values (U=16, p=.776), confirmed by a one-tailed Bayesian t-test (BF=.321). Visual inspection of the classification patterns, e.g. which finger is mistaken for which, hinted at reduced performance specifically for the index finger (D2). This was confirmed by a group comparison (U=6, p=.012) and strong support for the alternative hypothesis that D2 classification is lower in the anophthalmic individuals (BF=15.1). Together, these results make clear that finger selectivity in S1, as well as other measures of hand representation, are not increased in the anophthalmic individuals compared to sighted controls.

### Finger-specific information in the occipitotemporal cortex of anophthalmic individuals

Next, we examined finger representation in the occipitotemporal cortex, more specifically area hMT+. Finger movements (when contrasted against baseline) result in bilateral activity in area hMT+ in the anophthalmic individuals (see Figure 3A-C). In order to quantify the gross recruitment and any fine-scale organisation within hMT+, we calculated the measures introduced for S1 above in a pre-defined ROI (see Methods). Because we had no a priori predictions concerning laterality – cross-modal activity has previously been observed in either or both hemispheres – we analysed hMT+ for each hemisphere separately.

**Figure 3:**
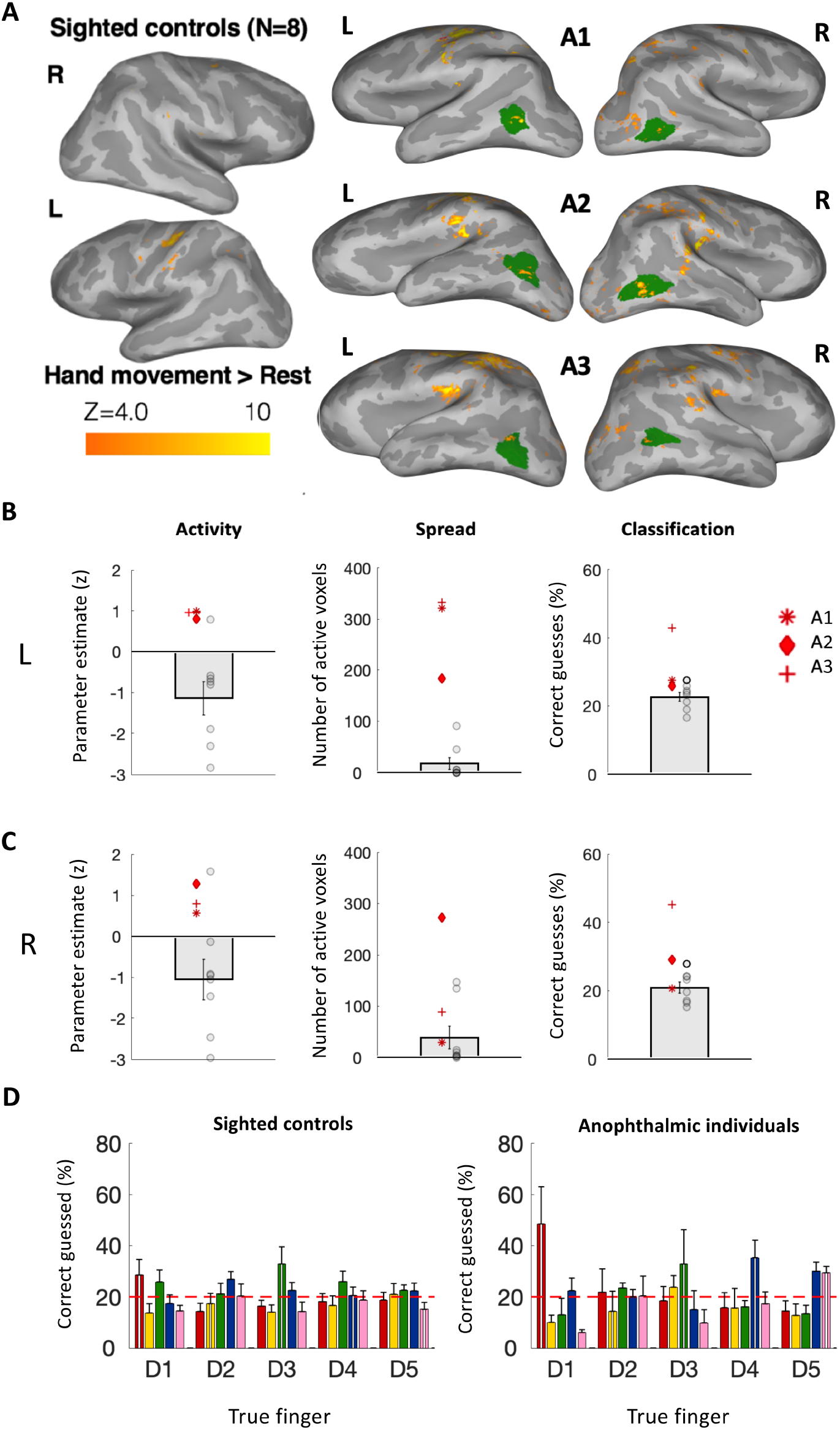
Hand representation in the human middle temporal complex (hMT+). **A)** Whole-brain activity for finger movement (contrasted against rest) in hMT+ for sighted controls (group average) and three anophthalmic individuals. Sighted controls do not show activity in hMT+, whereas the anophthalmic individuals do. Area hMT+ indicated in green. **B)** Proxy measures of hand representation in left hMT+. Activity and spread tended to be higher in anophthalmic individuals than sighted controls. Classification showed significant finger-specific information in all anophthalmic individuals and 1 control (indicated by bold circle). **C)** Proxy measures of hand representation in right hMT+. Activity and Spread group differences are trending, and classification showed significant finger-specific information in 2/3 anophthalmic individuals and 1 sighted control (indicated by bold circle). **D)** Average classification performance in left hMT+ across participants for sighted controls (left) and anophthalmic individuals (right). Note that the classifier performed best for finger D1 in the anophthalmic individuals. Other symbols as in Figure 2. L: left; R: right hemisphere.

In the left hemisphere, average activity was significantly greater for the blind group than for the sighted group (U=30, p=.012, BF=3.71). In the right hemisphere, this difference was not significant (U=27, p=0.085, BF=1.89). Spread of the activity showed similar lateralisation to the left (left hemisphere: U=30, p=.012, BF=959; right hemisphere: U=26, p=0.133, BF=1.85). All Bayesian t-tests had as their alternative hypothesis that anophthalmic individuals activate hMT+ differently than sighted controls (2-sided).

In order to determine whether the fine-scale finger organisation in hMT+ resembled that of S1, we initially correlated the representational structure (using RSA) to that that of S1. Yet, left hMT+ did not produce a reliable structure (indicated by poor split-half reliability; mean rho=.04). In fact, the split-half correlation was not even different from 0 (Wilcoxon signed rank test: rank=34; p=.966). Therefore, if any information content is facilitated by the increased activity, this is not reflected by a consistent representational structure measurable at the voxel level. Our hypothesis predicting that due to cross-modal plasticity, hMT+ would manifest an organisational structure similar to that of low-level somatosensory areas (greater overlap in representation across neighbouring fingers) was therefore not confirmed.

To examine digit information in hMT+ more directly, we repeated the classification analysis of individual fingers, as described for S1. In left hMT+, all anophthalmic individuals performed significantly above chance level (20%), whereas only 1 control performed better than chance. As a group, classification was significantly higher for the anophthalmic individuals (32%) than for the controls (23%; U=28.5, p=.036; BF=4.49; see Figure 3B). In right hMT+, classification performance was significant in 2/3 anophthalmic individual and 1 control.

Because not all anophthalmic individuals showed increased finger information, no follow-up group comparison was performed. To determine whether the increased information content in the left hemisphere was driven by specific fingers, misclassifications were examined separately per finger, for both groups (Figure 3C). If there were hidden topographic regularities in the finger representations, then the classifier should show more confusion between neighbouring fingers (as illustrated by the classification errors in the S1 topography, Figure 2D). Misclassifications did not seem to be structured in a certain way, except for the little finger (D5) often being mistaken for the ring finger (D4). Instead, the thumb (D1) was the most distinct of all classified fingers, in line with other literature (Ejaz et al. 2015). On the whole, improved (albeit slightly) classification performance in the anophthalmic individuals suggest there is more information about fingers, and the thumb in particular, in left hMT+ of anophthalmic individuals than of sighted controls.

Finally, as an exploratory confirmation that the increased finger information is at least specific to hMT+, we repeated the analysis in a visual control area V6/V6a. We found significant finger information in two sighted (left hemisphere), and a sighted and an anophthalmic participant (right hemisphere), suggesting some task-related information potentially exists in this area. As a group, in either hemisphere, classification was not significantly higher for the anophthalmic individuals (left: 22%, right: 23%) than for the controls (left: 25%; U=18, p=1; BF=1.85, right: 23%, U=17, p=.921, BF= 1.95). This analysis reaffirms the increased role of hMT+ in housing finger-specific information in the ‘visual’ cortex of anophthalmic individuals.

## Discussion

Here, we examined cortical representation of the hand in S1 and found that anophthalmic individuals did not show a more pronounced finger map than sighted controls, in any measure that was tested. Compared to sighted controls, there was, however, significantly more information about individual fingers in left hMT+ of the anophthalmic individuals.

### S1 hand organisation is stable in anophthalmic individuals

Previous studies reported that S1 hand representation of congenitally blind individuals is different from that of the sighted, in terms of spatial spread of activity (Pascual-Leone et al. 1993, Pascual-Leone and Torres 1993), strength of activity (Giriyappa et al. 2009), or a shifted inter-finger balance (Sterr et al. 1998). This has been interpreted as a positive adaptation towards more effective tactile processing for people who rely more on touch in their daily lives. Conversely, Gizewski et al. (2003) suggest there is little difference between blind and sighted participants in how the primary sensorimotor cortex is recruited during finger tapping. Also, despite frequently being cited as evidence for S1 plasticity, Burton et al. (2004) show no statistical differences in average S1 activity between early blind and sighted individuals following vibrotactile stimulation; although their data may show a weak effect of blindness on the spatial spread of said activity, this is not directly tested.

The study of sensorimotor behaviour in blind individuals is dominated by the study of Braille users. Similar to the debatable benefits of blindness for tactile performance (Norman and Bartholomew 2011, Alary et al. 2009, Wong et al. 2011), it is unclear how much impact Braille reading has. For example, grating orientation thresholds have been reported to be reduced on Braille reading fingers (Van Boven et al. 2000), but also not to be different between blind participants who do and do not read Braille (Norman and Bartholomew 2011). The role of Braille in everyday life of blind people has progressively been taken over by various forms of (non-tactile) technology (National Federation of the Blind Jernigan Institute 2009), exemplified by one of the anophthalmic individuals tested here who does not read Braille at all. The only differences with sighted controls we observed regarding S1 hand representation was poorer classification performance for the index finger (D2) of the anophthalmic individuals and greater spatial spread for D1 activity. The former finding seems to go against greater individuation of finger representation in blind individuals, which one would conjecture on the basis of more individuated finger usage during Braille reading (Mogilner et al. 1993, Vidyasagar et al. 2014). The latter also does not fit this theory, because our anophthalmic participants are not more prone to use their thumb than is typical.

Non-human evidence in favour of S1 changes following blindness, i.e. altered somatosensory receptive fields in the barrel cortex of blind rodents and cats (Rauschecker et al. 1992, Toldi et al. 1994), is also accompanied by altered behaviour (whisker use) from early life onwards. Yet, beyond Braille reading, blind individuals may actually not use their hands differently from sighted individuals, simply more. This may explain why the anophthalmic individuals’ S1 hand representation is not different from that of sighted controls. As a measure strongly correlated to everyday hand use (Ejaz et al. 2015), the canonical structure of hand representation has been found to be remarkably stable, despite hand loss (Kikkert et al. 2016, Wesselink et al. 2019), extensive behavioural training (Beukema et al. 2019), or expert musicianship (Ogawa et al. 2019). While average dissimilarity between fingers may vary, indicating altered signal strength, the representational structure is stable. Our anophthalmic individuals complement these findings by demonstrating both stable representational structure and dissimilarity. Although evidence exists that sensory cortex is capable of considerable reorganisation (congenital one-handers: Wesselink et al. 2019, expert foot users: Dempsey-Jones et al. 2019b), it appears that a large deviation from typical input early in life, rather than later, is paramount. Given a recent report of typical hand representation in two macaques who were near-blind in the first year of their life (Arcaro et al. 2019), it is not evident early life behaviour of blind individuals deviates far enough from normalcy to trigger representational change.

An alternative account is that the behaviour of blind individuals is sufficiently different to induce plasticity in the (adult) sensorimotor system, but that the plasticity only manifests transiently and depends heavily on the immediately preceding experience. Kolasinski et al. (2016b) showed altered hand representation after two fingers had been glued together for 24 hours, but it was not irreversibly altered. Similarly, only after a full day of Braille reading did Braille readers’ reading finger’s M1 representation demonstrate signs of expansion (Pascual-Leone et al. 1995); not after rest days without reading. Perhaps, previous support for sensorimotor plasticity following blindness was due to transient effects of immediate (Braille) experience, rather than blindness per se. This would be coherent with the S1 stability found in our anophthalmic participants. If so, some previous findings of sensorimotor plasticity (in blind individuals) may have been facilitated by short-term modulation of input gain [as e.g. seen during stimulation (Pleger et al. 2003)] instead of substantial reorganisation due to an increased dependence on touch.

### Sensorimotor information of individual fingers is increased in hMT+ of anophthalmic individuals

In our anophthalmic participants, we found increased activity in hMT+ compared to sighted controls, consistent with other studies showing this for a range of conditions causing blindness (see Burton 2003, Sathian and Stilla 2010 for review). Yet, increased activity does not necessarily underlie functionally meaningful processing, but may also be observed because of increased noise (e.g. due to reduced inhibition) or task irrelevant processing. As an indication of emergent functional processes, we interrogated, using multivariate analyses, whether finger-specific information is conveyed in this activity. Finger-specific information, measured by classification, was increased in the anophthalmic individuals, suggesting some emergent sensorimotor hand-related process in the occipitotemporal cortex (see Beauchamp et al. (2009) for further support of a lack of finger information in the visual cortex of the sighted). Because the activity was not structured similarly to that in low-level somatosensory areas (greater overlap in representation across neighbouring fingers), we suggest any emergent process(es) is/are not a low-level addition to the sensorimotor network.

What non-primary sensorimotor function could be supported by the increased activity in the anophthalmic individuals? Because a more accurate functional localiser (Huk, Dougherty & Heeger, 2002) cannot be run with blind participants, the large anatomically defined hMT+ used in the current study does not allow us to pinpoint the physiological origins of finger selectivity. Like in sighted participants, hMT+ has been shown to be responsive to auditory (Bedny, Konkle, Pelphrey, Saxe, & Pascual-Leone, 2010; Dormal, Rezk, Yakobov, Lepore, & Collignon, 2016; Huber, Jiang, & Fine, 2019; Jiang, Stecker, & Fine, 2014) and tactile motion (Matteau, Kupers, Ricciardi, Pietrini, & Ptito, 2010; Sani, et al., 2010) in groups of congenitally blind individuals. Our task, however, did not require any motion discrimination, either explicitly or implicitly. Alternatively, the finger-specific information could relate to a more general role of the occipitotemporal cortex, the region including and surrounding hMT+, in body representation. Previous research has suggested that the lateral occipitotemporal cortex surrounding motion-sensitive regions in sighted people (Weiner & Grill-Spector, 2011) may be involved in visual (Downing et al. 2001), sensorimotor (Astafiev et al. 2004, Orlov et al. 2010) and action related (Peelen and Downing 2005, Gallivan 2014) body-part representation. Although we and others (Beauchamp et al. 2009) did not find any indication of single-finger information in the sighted, despite using a sensitive classification method, other studies have reported some increased activity in sighted hMT+ during hand movement (Astafiev et al. 2004, Orlov et al. 2010, Amedi et al. 2002, Beauchamp et al. 2007). As such, finger information could potentially occur latently in sighted individuals and become unmasked in the absence of vision (Kupers and Ptito 2014).

In light of the stable finger representation in S1, are the changes in hMT+ sufficient to explain the improved tactile performance of the blind? We were unable to correlate measures of brain organisation to behaviour directly given our low sample size. Yet, regardless of how finely tuned low-level representations are, performance on a psychophysical task also depends on the readout of such representations by higher-order areas. For tactile discrimination, readout is likely done in higher-order areas outside of S1 (Romo and Salinas 2001, Stilla et al. 2007, Sathian et al. 2013) and can, like other decision-making tasks, be improved through training or experience. Notably, after minimal training, sighted controls may demonstrate similar tactile acuity to blind participants (Grant et al. 2000); see also (Wong et al. 2011). If sighted controls can show expert tactile acuity, a reorganised occipital cortex is unlikely to be a necessary contributor in anophthalmic individuals. Still, future research should study the higher-order somatosensory and frontal areas involved in readout. If hMT+ plays a role in tactile decision-making in blind individuals, these areas would be expected to receive inputs from hMT+.

In this study, we used a specialised patient group to ensure the occipitotemporal cortex has never received any visual input. In other forms of (congenital) blindness, some input, e.g. retinal waves, may still access and influence the visual cortex during development (Watkins et al. 2013). Most studies on cross-modal activity in diverse congenitally blind populations still show effects indicating cross-modal plasticity (Kupers and Ptito 2014), even in individuals who got blinded postnatally (typically within the first years; (Bedny 2017)). If anophthalmic individuals are not representative of the greater blind population, their idiosyncratic development likely restricts the scope for cross-modal plasticity less than other blind individuals; not more.

To conclude, we have studied three congenitally blind individuals with a rare condition that makes them particularly susceptible to cross-modal plasticity. Although they showed superior tactile acuity compared to sighted controls, the anophthalmic individuals showed no signs of stronger finger representation in S1, suggesting this area is organised similarly to that of sighted controls. In left hMT+, finger movements induced greater activity for the anophthalmic individuals than for controls. This activity did not express typical markers of low-level sensorimotor areas, but it contained finger-specific information. This suggest that area hMT+ can acquire some novel somatosensory processing that may not be typically present.

## Acknowledgements

TRM was funded by a Sir Henry Dale Fellowship jointly funded by the Wellcome Trust and the Royal Society (grant number 104128/Z/14/Z) and an ERC Starting Grant (grant number 0032-2-289-6121). For the purpose of Open Access, the author has applied a CC BY public copyright licence to any Author Accepted Manuscript (AAM) version arising from this submission. The authors would like to thank Samuel Hurley for extensive physics support during data collection and James Kolasinski for his comments on previous versions of this work.

## Material and methods

### Participants

Four individuals with bilateral Anophthalmia, a rare congenital condition in which both eyes fail to develop, participated in this experiment (hereafter, anophthalmic individuals; mean age: 33 ± 3; 3 male; 1 left-handed; see Table 1 for demographic details). One anophthalmic individual was excluded from all data analysis due to inability to perform the fMRI task (see fMRI task section below for details; this data is not reported in the current manuscript). The full experimental protocol involved a battery of behavioural tasks and fMRI measurements over the course of 1-2 sessions, as elaborated in the full protocol available online: https://osf.io/d9g3t/. Here, we focus on the procedures relevant to the key analyses described below. Eight sighted healthy control participants (mean age: 25 ± 3; three male) with normal (or corrected) vision underwent the same neuroimaging procedures. To compare fMRI S1 representation with a larger control group, further data was retrieved from a study using similar scanning parameters (n=15, 26 ± 4; 9 male; 1 left-handed; (Sanders et al. 2019)). One key difference with our current study protocol is that participants received visual feedback and were not blindfolded. All participants used their right hand for the various tasks.

**Table 1.** Demographic details. Demographic details of participants. A: Anophthalmic individuals; C: Sighted controls.

| ID Code | Age | Gender | Handedness | Braille hand(s) |
| --- | --- | --- | --- | --- |
| A1 | 36 | M | R | - |
| A2 | 31 | M | R | R & L |
| A3 | 30 | M | R | R & L |
| C1 | 24 | F | R | - |
| C2 | 22 | F | R | - |
| C3 | 27 | M | R | - |
| C4 | 26 | M | L | - |
| C5 | 22 | F | R | - |
| C6 | 30 | M | R | - |
| C7 | 25 | F | R | - |
| C8 | 25 | F | R | - |

In addition, to compare the tactile acuity of the anophthalmic individuals to a larger group of sighted control participants we used data collected as part of two larger studies: 1) For tactile acuity comparisons, data was retrieved from Dempsey-Jones et al. (2016; n=21, age=26.4±7.1, 1 left-handed) and [(Dempsey-Jones et al. 2019a); n=36, age=27.7±6.7, 5 lefthanded]. The anophthalmic tended to be older than the control group used for tactile acuity comparisons (U=143, p=.083). Previous research shows an advantage to younger participants in tactile acuity tasks (Legge et al. 2008). Therefore, the fact that the (older) anophthalmia participants outperformed their younger controls makes their better tactile performance even starker.

Experimental procedures were approved by the Medical Sciences Interdivisional Research Ethics Committee of the University of Oxford (C2-2013-05 and C1-2011-101). All participants gave their informed consent prior to the study.

### Behavioural task & analysis

Tactile acuity was measured for the index (D2), middle (D3) and ring finger (D4) of the right hand, using standard orientation grating testing. A detailed description of the task can be found in Dempsey-Jones et al. (2016). In short, participants were asked to make a tactile judgement on the perceived orientation (‘horizontal’ or ‘vertical’) of plastic dome gratings varying in groove width and isometric groove spacing (between .25 - 3.5 mm). Each grating was presented either parallel or orthogonally to the distal finger pad for 1-2 blocks of 10 trials on each finger. Blocks were randomly interspersed. At the end of each trial, participants reported the perceived orientation using a right or left button press on a mouse held in their left hand. No time limitations were posed, and participants were encouraged to perform the task as accurately as possible. A psychometric function was fit to the data using least squares fitting and the tactile threshold for each finger was interpolated at 82% accuracy (see Dempsey-Jones et al. 2016). Finally, the tactile thresholds were averaged across the three fingers for each participant.

### fMRI task

Finger representations were probed using an active finger-tapping task, adapted from Diedrichsen et al. (2013). This task has been previously shown to yield strong inter-and intra-subject reproducibility of SI finger representation [(Kolasinski et al. 2016a, Ejaz et al. 2015); see further validation against passive paradigms in (Berlot et al. 2019, Sanders et al. 2019)]. Finger movement recruits a combination of peripheral receptors, encoding a range of somatosensory modalities (e.g. proprioception and mechanoreception from surface and deeper receptors), as well as efferent information from the motor system. It is therefore more likely to engage the participants’ occipitotemporal cortex than passive finger stimulation. Specifically, our task involved individual finger movement blocks (8s each) for each of the five fingers, and a no movement (i.e. rest) condition (8-16s each). Each run was made up of four repetitions of each finger movement condition, presented in counter-balanced order, and five repeats of the rest condition at regular intervals. Example sequences are shown in Table S1. Each participant underwent 8 runs and run order was also counterbalanced across participants. Participants used a button box with four buttons to perform the task; they pressed the side of the box with their thumb. Participants were instructed to perform these button presses at a constant rate of ~1Hz, while keeping non-instructed fingers still.

Sighted participants were blindfolded and all participants were cued tactilely. The use of tactile cues was chosen over that of auditory ones to minimise the risk of engaging the tonotopic representations in the hMT+ in anophthalmic participants (Watkins et al, 2013). To cue the participant which finger should be moved, an experimenter lightly patted the participant’s leg at a specific location at the start of each movement block. The cues followed a spatial pattern that the participants were trained on prior to scanning: thumb movement was instructed by a touch at the ball of the left foot, index – outer left ankle, middle – inner left ankle, ring – inner right ankle, and little – outer right ankle. Rest blocks were cued by a double-tap on the balls of both feet simultaneously. The experimenter received auditory instructions specifying the block order through MR-compatible headphones. In order to ensure a steady movement pace of 1 Hz, participants were instructed to follow the sound of the vibrating magnetic coils of the MRI at every gradient switch. In our case, a gradient switch occurred every two seconds; participants were instructed to make two button presses between two gradient switches. To ensure good understanding of these instructions, the experimenter trained the participants on both the motor and the cueing component of the task. The auditory cueing to the experimenter and the recording of the button responses were implemented using Presentation software (version 0.70, www.neurobs.com). Correct task performance was further verified off-line using button box responses that were recorded during the experiment. Note that one anophthalmic individual was unable to maintain a steady 1Hz pace and was subsequently discarded from all analysis.

### MRI acquisition, pre-processing, and low level analysis

High-resolution BOLD fMRI data was obtained using a Siemens (Erlangen, Germany) whole-body 7 tesla Magnetom scanner with a 32-channel head coil. Task fMRI data was acquired using Multi-Band Echo Planer Imaging (EPI) with an acceleration factor of 2 (Moeller et al., 2010) and a limited field of view (FOV), designed to capture both the hand knob of the central sulcus and the lateral occipitotemporal cortex bilaterally. The following parameters were used: spatial resolution: 56 slices with a 192 x 192 in-plane FOV, 1mm isotropic spatial resolution, TR: 2000ms, TE: 25ms, PE acceleration factor: 3, flip angle: 85 °, and phase partial Fourier: 6/8. Please note that due to the increased spatial resolution and the consequential limited FOV, we were unable to acquire other potential areas of interest (e.g. secondary somatosensory cortex, cerebellum, primary visual cortex) consistently across participants. Fat suppression was done by CHESS. Because data acquisition expended a large part of the scanner’s CPU and memory capacity, image reconstruction was performed offline. For some participants this caused data acquisition delays or partially corrupted data. As a result, only 7, 6 and 4 usable runs were obtained for sighted control C2 and anophthalmic participants A1 and A3, respectively.

Anatomical T1-weighted (MPRAGE) whole-brain images with a 1 mm isotropic resolution was also acquired. When available, images were collected at using 3T MRI machine, as they suffer less from intensity biases due to field inhomogeneity (Vaughan et al., 2001). For all anophthalmic individuals and sighted participant P2, the anatomical T1-weighted image was acquired at 3T using standard parameters (TE: 6.0ms, TR: 15ms). For the remaining participants an anatomical T1-weighted image was acquired using the 7T MRI scanner in the same session as the fMRI measurements, using the following parameters: FA: 7°, TI: 1050ms; TE: 2.82ms; TR: 2200ms. Fat suppression was done by means of water excitation.

Data pre-processing, GLM analysis and cortical surface reconstruction were implemented using software from FSL (Smith et al. (2004); Jenkinson et al. (2012); fsl.fmrib.ox.ac.uk/fsl) and Freesurfer (Dale et al. (1999); www.freesurfer.net). Connectome Workbench software (www.humanconnectome.org) was used for visualisation on the cortical surface. These tools were complemented by scripts written in UNIX or Matlab (version 8.5, R2015a) which were developed in-house (https://github.com/ronimaimon).

Each individual imaging run was pre-processed using FEAT (version 6.0). This included the following steps: motion correction using MCFLIRT (Jenkinson et al. 2002), brain extraction using BET (Smith 2002), high-pass temporal filtering (100s), and spatial smoothing using a 2mm FWHM (full width at half maximum) Gaussian kernel. A medium level of smoothing was done to improve the sensitivity of our multivariate analysis, while preserving spatial specificity (Hendriks et al. 2017). In addition, the motion estimates were inspected for excessive participant motion: no run contained more than 1mm (i.e. our voxel size) of relative displacement.

Images were registered to each other using FLIRT (Jenkinson et al. 2002). First, all fMRI runs were realigned to their average space (midspace), by iteratively averaging and coregistering. This procedure ensured the final coregistration was minimally biased towards a single run and all runs were minimally reoriented. Then, all functional images were registered to the anatomical volume. All coregistration was visually checked and manually adjusted if needed. Anatomical T1 images were further used to reconstruct the pial and grey matter inflated surfaces using Freesurfer (http://surfer.nmr.mgh.harvard.edu).

First-level parameter estimates were computed for each run using a voxel-wise general linear model (GLM), as implemented in FEAT, based on the double-gamma hemodynamic response function (HRF) and its temporal derivative. 11 contrasts were defined: each individual finger movement versus rest, movement of all fingers versus rest, and each individual finger movement versus movement of all other fingers. The voxel-wise parameter estimates were averaged across runs for each participant using a fixed effects model. The resulting Z-statistic images were thresholded using clusters determined by Z > 3.1 and p < 0 .05 family-wise-error-corrected cluster significance thresholding was applied. For visualisation purposes, the resulting Z-statistic images were projected to two-dimensional surface space using a cortical ribbon mapping method.

### Regions of interest

Regions of interest (ROIs) were defined on the cortical surface. To initially identify the anatomical hand regions in S1, a rectangle was drawn on the Freesurfer standard surface stretching 2 cm medially and 2 cm laterally to the anatomical location of the hand knob (Yousry et al. 1997) within areas BA3a, BA3b and BA1, defined by Freesurfer based on ex-vivo cytoarchitectural data. This is where finger maps are typically reported. This anatomical hand ROI was then projected to each individual surface. Next, all voxels within said rectangle that showed selectivity for at least one finger (i.e. contrast any one finger versus the others, Z>3.1) were grouped together for each participant to create the final individualised hand ROI. By choosing finger-selective voxels, we improved the signal-to-noise ratio, necessary because of the single-case inferences. Importantly, since the size of the ROI was not different between groups, any biases due to ROI selection should not impact the results.

To delineate the regions of interest within the visual cortex, each individual anatomical surface was aligned to the cortical atlas of the Human Connectome Project (Glasser et al. 2016). ROIs for the functional region hMT+ was created by combining regions FST, V4t, MT and MST. Various definitions exist for hMT+ and our liberal anatomical definition should overlap with previously functionally defined ROIs for hMT+ (Dumoulin et al. 2000). since our main methods (representational similarity analysis and classification) can cope with unnecessary non-informative voxels, this was preferable to failing to capture any emergent representation that is just outside MT/MST proper. This methodological choice has likely influenced the relatively low average activity reported in Figure 3B. ROIs were defined separately for the left and right hemisphere.

### fMRI analysis

#### Univariate analysis

In order to visualise finger maps in S1, for each participant the Z-statistic images for each contrast “1 finger > mean of other fingers” were averaged across runs and projected onto the participant’s inflated surface. Univariate inference was done on two measures: 1) average activity (mean all fingers versus rest) inside the ROI, and; 2) number of active voxels (mean of all fingers > rest; Z > 3.1) inside the ROI, corrected for whole-brain false discovery rate (q(FDR) = .01).

#### Representational similarity analysis

In order to assess whether hand-related activity contained a representational structure within an ROI, the dissimilarity in activity patterns (for each finger versus rest) were calculated for each finger pair for each participant and ROI. Our dissimilarity measure was the Mahalanobis distance (Nili et al., 2014; Ejaz et al., 2015), cross-validated over each possible pair of runs and then averaged. This ensures that the expected distance is zero if two patterns are not statistically different from each other, whereas larger distance values indicate greater dissimilarity/separation. The resultant inter-finger distances were arranged into one dissimilarity matrix (RDM) per participant per ROI. Next, we extracted the mean dissimilarity between all possible finger pairs as an indication of finger separability. If there is no representational structure, one would expect the average dissimilarity (or separability) to be 0. We further calculated the Spearman correlation between the participant specific RDMs and the sighted controls’ average RDM of the S1 hand area (using a leave-one-out approach for the controls). In order to assess whether RDMs were reliable at the level of the individual participant, this process was repeated separately for odd and even runs. Decent cross-half correlation (rho>.6; a liberal threshold: (Kline 2000)) between ‘odd and even RDMs’ was a prerequisite to interpret the RDM calculated using all runs.

#### Classification

Classification was implemented using the Princeton MVPA toolbox for Matlab available at https://code.google.com/p/princeton-mvpa-toolbox/ (see Wolbers et al. (2011) for an implemented example). The classifier algorithm used consisted of a backpropagation neural network with one hidden layer consisting of 10 nodes (Duda et al., 2012). In short, the network was trained to classify activity patterns for each finger and subsequently had to classify unseen patterns as belonging to 1 of the 5 tested fingers. The classification was crossvalidated in a leave-1-run-out fashion. For example, training was performed on 7 runs x 5 finger patterns and testing on the remaining 5 finger patterns from 1 imaging run, which was then repeated such that each run functioned as a test once. For each ROI, this classification was repeated 20 times with random initial weights and averaged, in order to minimise the effect of the network’s initial state. Final classification accuracy was calculated by averaging across the 5 individual fingers. Given the 5 fingers on a hand, chance level was 20%. In addition, we wanted to determine whether the classification performance was significant in individual participants. For this purpose, classification performance was compared against a chance distribution simulated for each ROI and participant. This distribution was created by running the above classification on the same data but with finger identify scrambled within each run. This was repeated 200 times and significance was determined by a one-sided ranked-order test.

#### Statistical analysis

Although it is typical for groups with a low sample size to be studied as multiple single-case tests, we report mainly group comparisons. In our exploration of S1, this made it more likely to find any differences. Single-case inference led to inconclusive findings (Crawford and Garthwaite 2007). In the visual cortex, differences between the sighted group and anophthalmic participants have been tested individually. All group comparisons were done using Mann-Whitney U tests (α = 0.05), suitable for small groups, as implemented in Matlab. In order to assess our confidence in supporting any null results, we used Bayesian t-tests, as implemented in JASP (https://jasp-stats.org/). The specific alternative hypothesis that was tested (greater than, smaller than, or different from the null), is specified in the main text for each test. A Bayes Factor (BF) above 3 was interpreted as substantial evidence for the alternative hypothesis; a BF below 0.33 as substantial evidence towards the null (Kass and Raftery 1995).

---

**Supplementary Table S1.**
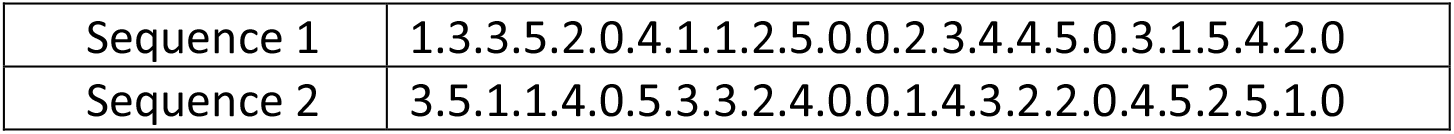
Example movement sequences from the active movement task. Numbers indicate a block of 8 trials consisting of one finger (thumb-little finger; 1-5) moving, or 8 seconds of rest (0). Each sequence was counter-balanced; the whole set was counter-balanced for 1-back sequences.

|  |  |
| --- | --- |
| Sequence 1 | 1.3.3.5.2.0.4.1.1.2.5.0.0.2.3.4.4.5.0.3.1.5.4.2.0 |
| Sequence 2 | 3.5.1.1.4.0.5.3.3.2.4.0.0.1.4.3.2.2.0.4.5.2.5.1.0 |

